# Role of NUMTs (Nuclear mitochondrial DNA) genes in affecting disease resistance in duck against Pasteurellosis

**DOI:** 10.1101/2023.08.30.555528

**Authors:** Jyoti Sahu, Aruna Pal, Argha Chakraborty, Samiddha Banerjee, Manti Debnath, Rajarshi Samanta

## Abstract

Ducks are mostly resistant to common avian diseases, but frequent occurrence of duck pasteurellosis, commonly kown as Duck Cholera, caused by *Pasteurella multocida* may cause a loss. In our earlier studies, we have identified certain immune response genes of nuclear origin as well as mitochondrial genes in duck, conferring resistance against duck cholera. In our current study, we have detected certain NUMT (Nuclear mitochondrial) genes in duck with certain role in disease resistance in case of duck cholera. NUMT genes are basically nuclear genes, but they act through mitochondria. Identified NUMT genes (Thymidine phosphorylase/ endothelial cell growth factor1 gene, TFAM Transcription factor A, mitochondrial, TK2 Thymidine kinase 2) were characterized and certain important domains were identified. Differential mRNA expression profiling revealed upregulation of the genes in healthy ducks compared to that of infected ducks. Exploitation of the identified genes may lead to development of ducks resistant to duck Cholera.

## Introduction

Ducks have been considered to be the second most important poultry species after chicken. From historic times, ducks have been reared domestically for food purpose. In recent years this duck farming has been growing gradually and is becoming a remarkable segment in poultry meat industry especially in Asia. The duck industry has commenced to follow the same pattern of broiler industry with the increasing demand. India stands 8^th^ in total duck production in world. According to 20th Livestock Census (2019) in India total duck population was 33.511 million with 32.50 million in backyard duck farming, and 1.08 million in commercial poultry farm. The improved duck and desi duck contribute 0.21% and 0.86%respectively with respect to total egg production (Livestock Census,2019).

Indigenous ducks have been observed to have certain unique characteristics in terms of its disease resistance ability against common avian diseases (Pal et al., 2021). Ducks can be easily reared under organized farming system with small water container that enable them to dip their head for washing of their eyes without any natural water bodies. Ducks grow faster than chicken. We had already conducted certain preliminary studies on duck (Pal, 2020, Debnath et al, 2022). They have high prolificacy than chicken. However Duck cholera or Duck pasteurellosis is a challenging disease impacting economic loss.Duck pasteurellosis caused by Pasteurella multocida may occur as acute, chronic or in occassional cases, it may even be asymptomatic (Pal et al., 2021). Pasteurella multocida is a gram negative facultative commensal bacteria. It has been classified as serotypes 1 to 16 as per their lipopolysaccharide antigen and serogroups A, B, D, E and F based on its capsular component. Serogroup A are generally responsible for fowl cholera. In the early stages of an illness, mortality rates might range from 5% to 20%. Even far higher rates of mortality could occur (Pilatti et al., 2016). Mortality may decrease to 2%–5% each month as the illness progresses into chronicity. Birds with a chronic infection may pass away, suffer from infection for a long time, or recover. An outbreak flock of 50,000 birds had an average cost of $0.40 (€0.33) per bird, while nonoutbreak flocks that had received the fowl cholera vaccine had an average cost of $0.12 (€0.10) per bird. Furthermore, Christiansen (1990) discovered that the cost of treating an outbreak of fowl cholera in 1985–86 was estimated to be $0.51 (€0.452) per bird. This cost included productivity losses as well. About half as much was spent on the immunisation against poultry cholera overall as there was on the outbreak. In all, California lost 0.5% of its whole 1985–1986 turkey meat supply due to fowl cholera3.

In our lab, we had been studying on disease genetics in sheep(ref) and duck model (ref). We have studied immunogenetics against parasites, viral infection as duck plague and avian influenza (). In this study, we are working on host immunogenetics against bacterial infection. We had studied immunogenetics based on nulcear genes(ref) and mitochondrial genes (ref). But in this current study we aim to study a novel class of gene, which are basically nuclear genes, but act through mitochondria, known as Nuclear mitochondrial DNA or NUMTs. Promising NUMT genes are thymidine phosphorylase (TP), Transcription factor A, mitochondrial (TFAM), Thymidine kinase 2, mitochondrial (TK2).

TYMP/ECGF1 encodes for the protein thymidine phosphorylase (TP), also known as TdRPase, or endothelial cell growth factor1, Gliostatin and Platelet-derived endothelial cell growth factor (PD-ECGF) (Akiyama *et al*., 2004, NCBI, 2022 Gene ID: 1890). In mice, it is comprised of 471 amino acids, in rat 476 amino acids and 482 amino acids in humans (Uniprot, P19971 • TYPH_HUMAN, Uniprot, Q99N42 • TYPH_MOUSE Uniprot, Q5FVR2 • TYPH_RAT). Nishino *et al.,* (1999) revealed its ***involvement in mitochondrial genome maintenance***, regulation of gastric motility, regulation of myelination and regulation of transmission of nerve impulses. Accordingly, this gene was observed to be involved in the disease of Mitochondrial DNA depletion syndrome 1, MNGIE type (MTDPS1) in humans which is a multisystem disease associated with mitochondrial dysfunction (Nishino *et al.,* 1999, Gamez *et al.,* 2002). This disease is clinically characterized by its onset between the second and fifth decades of life; there is ptosis, gastrointestinal dysmotility (often pseudoobstruction), progressive external ophthalmoplegia, cachexia, diffuse leukoencephalopathy, peripheral neuropathy and myopathy.

**<u>TFAM</u>** codes 246 amino acids, encodes for protein **Transcription factor A, mitochondrial**. Celestini *et al.,* (2018) and Hao *et al.,* (2020) reported that this gene binds to the mitochondrial light strand promoter and functions in mitochondrial transcription regulation. This gene was observed to be the component of the mitochondrial transcription initiation complex, composed at least of TFB2M, TFAM and POLRMT that is required for basal transcription of mitochondrial DNA (Hillen *et al.,* 2017). It was also reported the involvement of this gene in transcription initiation at the mitochondrial promoter and positively regulates DNA-templated transcription (Litonin *et al.,* 2010). This gene is required for accurate and efficient promoter recognition by the mitochondrial RNA polymerase. It promotes transcription initiation from the HSP1 and the light strand promoter by binding immediately upstream of transcriptional start sites and unwinding of DNA (Rubio-Cosials *et al.,* 2011). TFAM knockdown causes a reduction of mtDNA copy number and abnormalities in the embryos of Zebrafish (Otten *et al*., 2020). TFAM was observed to bend the mitochondrial light strand promoter DNA into a U-turn shape via its HMG boxes (Fisher, 1992). It is also required for the maintenance of normal levels of mitochondrial DNA (Gangelhoff *et al.,* 2009 and Kasashima *et al.,* 2012) as well as also plays a role in organizing and compacting mitochondrial DNA enabling chromatin binding (Hao *et al.,* 2020, Celestini *et al.,* 2018, Hillen *et al.,* 2017, Kasashima *et al.,* 2012, Rubio-Cosials *et al.,* 2011, Ngo *et al.,* 2011, Litonin *et al.,* 2010, Gangelhoff *et al.,* 2009, Fisher, 1992). TFAM was observed to be a part of a protein-containing complex, which is located in the mitochondrial nucleoid (Bogenhagen *et al.,* 2008, Celestini *et al.,* 2018). Gangelhoff *et al.,* (2009) reported this gene enables mitochondrial promoter sequence-specific DNA binding. TFAM is involved in mitochondrial transcription and enables mitochondrial transcription factor activity (Ngo *et al.,* 2011). Gaudet *et al.,* (2011) reported this gene enables transcription cis-regulatory region binding and DNA binding and bending. He also revealed its involvement in mitochondrial respiratory chain complex assembly. Due to mutation in this gene, a severe disease condition is seen known as Mitochondrial DNA depletion syndrome 15, hepatocerebral type (MTDPS15), which is characterized by severe intrauterine growth restriction, neonatal-onset hypoglycemia and liver dysfunction, there is also depletion of mitochondrial DNA in the liver and skeletal muscle and abnormal mitochondrial morphology is observed in skeletal muscle (Stiles *et al.,* 2016).

**<u>TK2</u>** encodes for the protein **Thymidine kinase 2, mitochondrial**. It is composed of 270 amino acids in *Mus musculus* and 265 amino acids in *Homo sapiens.* It was reported that TK2 phosphorylates thymidine, deoxycytidine and deoxyuridine in the mitochondrial matrix (Wang and Eriksson 2000, Wang *et al.,* 1999 and Saada *et al.,* 2001). They also reported in non-replicating cells, where cytosolic dNTP synthesis is down-regulated, mtDNA synthesis depends solely on TK2 and DGUOK. This gene can be widely used as a target of antiviral and chemotherapeutic agents. They also reported it enables deoxycytidine kinase activity. Gaudet *et al.,* (2011) reported this gene is active in the cytoplasm and enables deoxynucleoside kinase activity. TK2 gene is also active in the mitochondrion enabling thymidine kinase activity (Wang *et al.,* 1999, Saada *et al.,* 2001, Wang and Eriksson, 2000). Akman *et al.,* (2008) reported its involvement in the mitochondrial DNA metabolic process and mitochondrial DNA replication (Zhou *et al.,* 2008). This gene is also involved in the deoxycytidine metabolic process, DNA biosynthetic process, phosphorylation, thymidine metabolic process, nucleobase-containing compound metabolic process and pyrimidine nucleoside salvage. Mutation in the Thymidine kinase 2 gene causes the disease “Mitochondrial DNA depletion syndrome 2 (MTDPS2)”, which is characterized by the childhood onset of muscle weakness associated with the depletion of mtDNA in skeletal muscle. There is wide clinical variability; some patients have onset in infancy and show a rapidly progressive course with early death due to respiratory failure, whereas others have a later onset of a slowly progressive myopathy in humans (Saada *et al.,* 2001, Mancuso *et al.,* 2002, Wang *et al.,* 2005, Tulinius *et al.,* 2005 and Knierim *et al.,* 2015).

The previous works on NUMT genes has been done on mainly human or insect models. So the objectives are to explore the effects of these gene in indigenous ducks based on following considerations: Characterization of NUMT genes in indigenous duck affecting disease resistance against Duck Pasteurellosis and Differential mRNA expression profiling for the identified NUMT genes affecting disease resistance against Duck Pasteurellosis.

## Materials and Methods

### Characterization of Thymidine phosphorylase/ endothelial cell growth factor1 gene, TFAM Transcription factor A, mitochondrial, TK2 Thymidine kinase 2 in duck

#### Birds and sample collection

The samples were collected from Bengal ducks maintained at the West Bengal University of Animal and Fishery Sciences. Tissue samples were also collected from the slaughter house for Bengal ducks.

#### Sample Collection and RNA Isolation

Duck liver and kidney tissue tissue (1g) were collected from slaughtered birds which were clinically healthy and maintained in the duck house of the LFC Dept, WBUAFS. Adult males (n=6) were selected for the collection of samples. The tissue was immersed in Trizol in the vial and transported in ice to the laboratory for RNA isolation. Total RNA was isolated using the TRIzol extraction method (Life Technologies, USA), as per the standard procedure and was further utilized for cDNA synthesis (Pal and Chatterjee, 2009; Pal et al., 2011, Banerjee et al., 2021, Rawat et al., 2021). cDNA concentration was estimated and samples above 1200 micrograms per ml were considered for further study.

#### cDNA synthesis and PCR Amplification of TYMP/ECGF1, TFAM, TK2 Gene of Bengal Duck

20μL of the reaction mixture was composed of 5μg of total RNA, 0.5μg of oligo dT primer (16–18mer), 40U of Ribonuclease inhibitor, 1000M of dNTP mix, 10mM of DTT, and 5U of MuMLV reverse transcriptase in an appropriate buffer. The reaction mixture was mixed thoroughly followed by incubation at 37°C for 1 hour. The reaction was allowed up to 10 minutes by heating the mixture at 70°C unliganded and then chilled on ice. Afterwards, the integrity of the cDNA was checked by performing PCR (Pal, A, 2021, Pal and Chatterjee, 2010, Pal et al., 2014a,b, Pal et al., 2019ab, Pal et al., 2020, Pal et 2011). The concentration of cDNA was estimated through Nanodrop. TYMP/ECGF1, TFAM, TK2 gene primer pairs were designed based on the published sequences of chicken and duck origin using Primer 3 plus software (https://www.primer3plus.com) to amplify full-length open reading frame (ORF) of these genes (Table 1).

**Table 1a:**
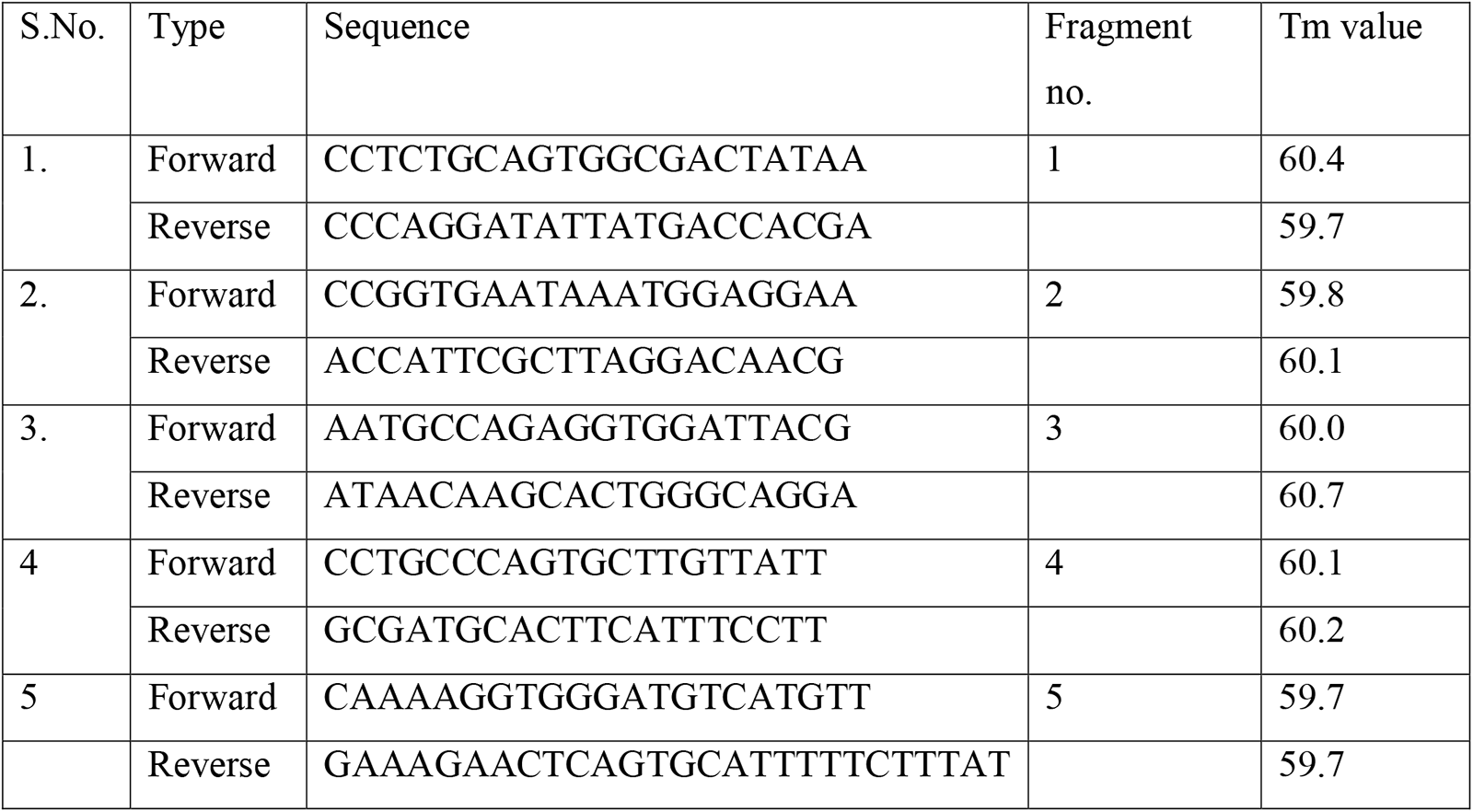
Primer sequences for Thymidine kinase (TK)

**Table 1b:**
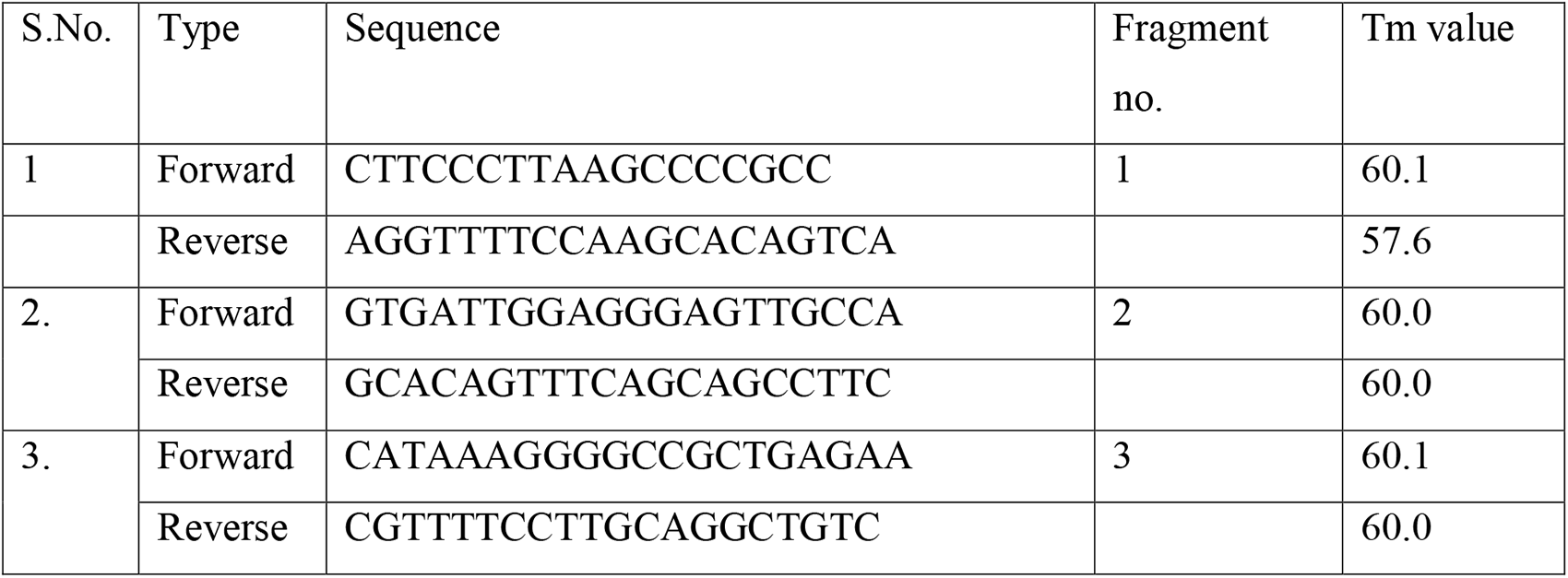
Primer sequences for TFAM Transcription factor A, mitochondrial for duck.

**Table 1c:**
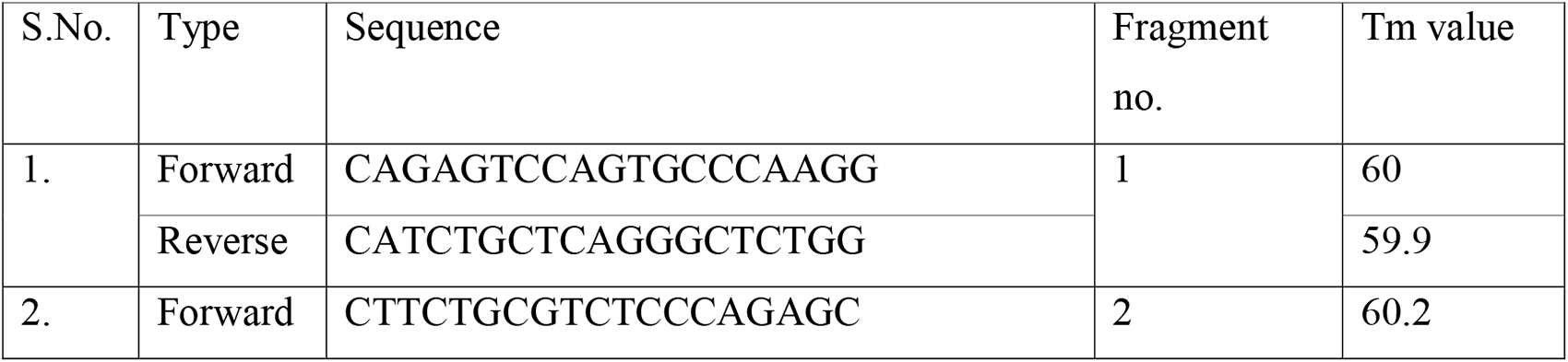
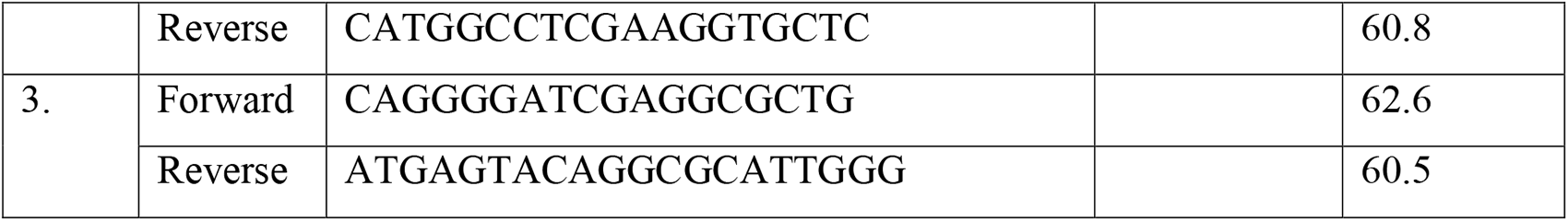
Primer sequences for *Thymidine phosphorylase/ endothelial cell growth factor1 gene* TYMP/ECGF1.

25μL of reaction mixture was comprised of 80–100ng cDNA, 3.0μL 10X PCR assay buffer, 0.5μL of 10mM dNTP, 1U Taq DNA polymerase, 60ng of each primer, and 2mM MgCl2. PCR reactions were performed in a thermocycler (PTC-200, MJ Research, USA) with cycling conditions as, initial denaturation at 94°C for 3min, further denaturation at 94°C for 30sec, annealing at 61°C for 35sec, and extension at 72°C for 3min were conducted for 35 cycles followed by a final extension at 72°C for 10 min.

#### cDNA Cloning and Sequencing

Amplified products of TK, TFAM, TYMP were checked by 1% agarose gel electrophoresis. The products were purified from gel using a Gel extraction kit (Qiagen GmbH, Hilden, Germany) .pGEM-T easy cloning vector (Promega, Madison, WI, USA) was used for cloning. Then, 10μL of the ligated product was mixed thoroughly with 200μL competent cells, and heat shock was given at 42°C for 45sec in a water bath. Subsequently, the cells were immediately transferred on chilled ice for 5 min., and SOC medium was added to it. The bacterial culture was centrifuged to obtain the pellet and plated on an LB agar plate containing Ampicillin (100mg/mL) added to the agar plate @1:1000, IPTG (200mg/mL) and X-Gal (20mg/mL) for blue-white screening. Plasmid isolation from overnight-grown culture was carried out by a small-scale alkaline lysis method as described in (Pal et al., 2021, Sambrook et al., 2001). Recombinant plasmids were characterized by PCR using primers as reported earlier and restriction enzyme digestion. The gene fragments released by enzyme EcoRI (MBI Fermentas, USA) were inserted in a recombinant plasmid which was sequenced by the dideoxy chain termination method with T7 and SP6 primers in an automated sequencer (ABI prism, Chromous Biotech, Bangalore).

#### Sequence Analysis

DNASTAR Version 4.0, Inc., USA software was employed for the nucleotide sequence analysis for protein translation, sequence alignments, and contigs comparisons (Pal et al.,2004, Pal et al., 2005). Novel sequences were submitted to the NCBI Genbank and accession numbers were obtained which are available in the public domain.

#### Study of Predicted duck TYMP/ECGF1, TFAM, TK2 Protein Using Bioinformatics Tools

Predicted peptide sequences of duck TYMP/ECGF1, TFAM, TK2 Protein were then aligned with that of other species using MAFFT (Katoh et al., 2013). The analysis was conducted for a sequence-based comparative study. The signal peptide is essential to prompt a cell to translocate the protein, usually to the cellular membrane and ultimately signal peptide is cleaved to give a mature protein. Prediction of the presence and location of the signal peptide of these genes was conducted using the software (SignalP 3.0 Sewer-prediction results, Technical University of Denmark). The leucine percentage was calculated manually from the predicted peptide sequence. Di-sulphide bonds are essential for protein folding and stability, ultimately. It is the 3D structure of the protein which is biologically active.

Protein sequence level analysis was employed (http://www.expasy.org./tools/blast/) for the assessment of leucine-rich repeats (LRR), leucine zipper, detection of Leucine-rich nuclear export signals (NES), and detection of the position of GPI anchor, N-linked glycosylation sites. Since these are receptors, they are rich in leucine-rich repeats, which are essential for pathogen recognition and binding. A leucine zipper is essential to assess the dimerization of IR molecules. N-linked glycosylation is important for the molecule to determine its membranous or soluble form. The leucine-rich nuclear export signal is essential for the export of this protein from the nucleus to the cytoplasm, whereas GPI anchor is responsible for anchoring in the case of membranous protein. Prosite was used for LRR site detection.

Leucine-rich nuclear export signals (NES) were analyzed with NetNES 1.1 Server, Technical University of Denmark. O-linked glycosylation sites were detected using NetOGlyc 3.1 server (http://www.expassy.org/), whereas N-linked glycosylation sites were assessed through NetNGlyc 1.0 software (http://www.expassy.org/). Sites for leucine-zipper were detected through Expassy software, Technical University of Denmark (Glick, 1977). Sites for alpha helix and beta sheet were detected using NetSurfP-Protein Surface Accessibility and Secondary Structure Predictions, Technical University of Denmark (Petersen et al., 2009). Domain linker sites were predicted (Ebina et al., 2009). LPS-binding (Cunningham et al., 2000) and LPS-signalling sites (Muroi et al., 2002) were predicted based on homology studies with other species of respective polypeptide. These sites are important for pathogen recognition and binding.

#### Three-dimensional structure prediction and Model quality assessment

Three-dimensional models of TYMP/ECGF1, TFAM, TK2 polypeptide were predicted through the Swissmodel repository (Kiefer et al., 2009). Templates possessing the greatest identity of sequences with our target template were identified with PSI-BLAST (http://blast.ncbi.nlm.nih.gov/Blast). PHYRE2 server based on the ‘Homology modelling approach was used to build three dimensional model of these proteins (Kelley, 2015). Molecular visualization tool as PyMOL (http://www.pymol.org/) was employed for model generation and visualization of the three-dimensional structure of these proteins understudies for duck origin. The structure of duck molecules was evaluated and assessed for its stereochemical quality (through SAVES, Structural Analysis and Verification Server, http://nihserver.mbi.ucla.edu/SAVES/); then refined and validated through ProSA (Protein Structure Analysis) web server (https://prosa.services.came.sbg.ac.at/prosa) (Wiederstein and Sippl, 2007). NetSurfP server (http://www.cbs.dtu.dk/services/NetSurfP/ Peterson et al., 2009) was used for assessing the surface area of these proteins through relative surface accessibility, Z-fit score, and probability of alpha-Helix, beta-strand and coil score.

The alignment of 3-D structure of these proteins was analyzed with RMSD estimation to evaluate the structural differentiation by TM-align software (Zhang et al., 2005).

#### Protein-protein interaction network depiction

To understand the protein interaction network of these proteins, we performed the search in STRING 9.1 database (Franceschini et al., 2015). The functional interaction was assessed with a confidence score. Interactions with scores < 0.3, scores ranging from 0.3 to 0.7, and scores >0.7 are classified as low, medium and high confidence respectively. Also, we executed a KEGG analysis which depicted the functional association of these proteins with other related proteins.

### Challenge study with Pasteurella multocida, symptomatic diagnosis, estimation of infective viral load at different stages of infection, PCR based detection

For the challenge study, initially, we incubated the fertile duck eggs (n=12) in the incubator at 37^0^C along with water in a Petri dish to maintain the proper humidity, which is essential for the incubation of duck eggs. After one week, we rechecked and screened the fertile embryo through candling. To harvest the *Pasteurella multocida* bacterial culture, at nine days of incubation of the fertilized egg, we puncture the air sac with the help of a sterilized needle through the chorioallantoic membrane and inoculated 200ul of the infective agent in a sterile environment. The needle puncture was quickly sealed with glue (feviquick). The eggs were again incubated in a sterile environment at 37C and humid conditions as earlier. The entire procedure was undertaken at the BSL3 lab. The duck plague viral strain employed was the available infection in the state from the clinical infected cases. After 3 days of inoculation, the viruses were isolated from embryonic fibroblast cells through a viral DNA isolation kit. Quantification of viral load was estimated through quantitative PCR with duck plague vaccine as control. To challenge, 0.3ml of viral stock at EID50 (Egg infective dose 50) was employed for each bird I/M. Infective amnio-allantoic fluid was used to determine 50 per cent EID. Initial verification was conducted to assure that the ducks were free from DP viral infection.

#### Sample Collection

For the present study, six tissue samples were collected from the infected indigenous duck samples followed by a challenge study. We designate the ducks with mortality as infected or diseased (less immune) and the ducks that survived as healthy (better immune).

#### Diagnosis and confirmation of Pasteurella multocida

The ducks were initially diagnosed with duck plague clinically with the help of initial symptoms. Later on, the confirmatory diagnosis was conducted through molecular detection of the Duck plague virus in the infected samples.

#### Gross Anatomical view of the organs in Pasteurella infected duck

We studied different body organs from infected ducks for clinical diagnosis of duck plague.

#### Confirmation through molecular detection

In the next step, we follow the confirmation of the samples with a molecular PCR-based technique using the following primers as recommended by OIE, Paris.

**Table 2:**
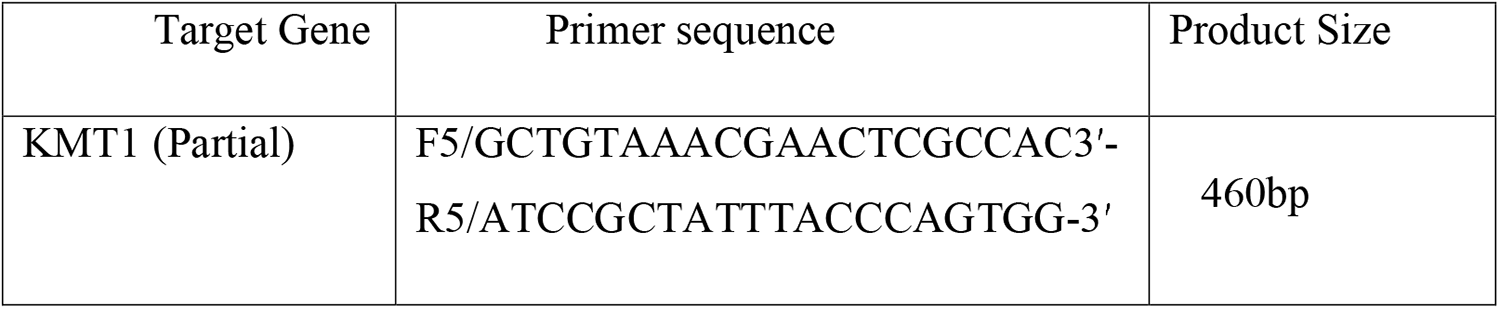
Primers used for the molecular detection of **Pasteurella multocida**.

It includes bacterial DNA isolation through an isolation kit and is subjected to PCR.

#### Assessment of dose of infection

We estimated the dose of bacterial infection standardization through Quantitative PCR. We take the standard vaccine as a control for its amplification and quantification.

#### Challenge study with Pasteurella multocida and study of expression profiling in healthy and infective groups

Differential mRNA expression profiling for GALT tissues w.r.t. **TYMP/ECGF1, TFAM, TK2** gene.

#### RT-PCR (Real Time-Polymerase Chain Reaction)

##### RNA Estimation

The tissue samples were cut into small pieces and submerged in liquid Trizol followed by thorough grinding with a mortar and pestle. 1 ml triturated tissue was taken into a 2 ml Eppendorf tube and 1 ml chloroform was added. Then it was centrifuged at the rate of 10,000 rpm for 10 minutes at 4°C in an automated temperature-controlled refrigerated centrifuge machine (BR Biochem, Life Sciences and REMI C24 plus). Three phases of differentiation were identified. The uppermost aqueous phase was collected for RNA isolation into a new Eppendorf tube and an equal volume of 100% isopropanol was added. It was left for a minute at 20-25°C temperature and centrifuged at the rate of 10,000 rpm for 10 minutes. After that supernatant was discarded and an equal volume of 70% ethanol was added to the pellet. Again the mixture was centrifuged at the rate of 10,000 rpm for 10 minutes and then the supernatant was discarded. Then the Eppendorf tube containing the pellet was kept at room temperature for air drying. After drying the remaining pellet dissolved with nuclease-free water (Applied Bio-systems, Cat. No. AM9930) and stored at -20°C for future uses (Pal et al., 2011, Pal, 2009, Pal et al., 2014).

##### Qualitative Analysis of Total RNA

The total RNA was quantified using NanoDrop-8000 (Thermo Scientific, Model No. 8000 Spectrophotometer), taking 1µl of each sample to determine the concentration and A260/280 ratio (Rawat et al. 2021, Banerjee et al., 2021, Pradhan et al., 2018).

##### First Strand cDNA Synthesis (Reverse Transcriptase -PCR)

5µl estimated RNA was taken into a PCR micro tube and added 1 µl OligoDT primer, 1µl 10mM dNTP and 13µl distilled water. After well mixing, it was heated at 65°C for 5 minutes. Then those tubes were quickly chilled on ice for 5-7 minutes and centrifuged. Then 4 µl 5x first strand buffer and 2 µl DTT (100mM) were added to the tube. It was well mixed and incubated at 37°C for 2 minutes. Then 1 µl M-MLVRT (200 u/µl) was added to the reaction mixture and treated at 37°C for 50 minutes and at 70°C for 15 minutes for inactivation. All the temperature treatments were maintained automatically in the thermal cycler machine (Applied Biosystems by Life Technologies) by setting the whole programme.

##### Primer Designing and Synthesis

All the primers were designed using primer 3 software (v. 0. 4.0) as per the recommended criteria. The primers for the genes used in the experiment, were designed (DNASTAR software).

##### Real-Time PCR Experiment (SYBR Green based)

The primers were standardised with respective cDNA samples before being subjected to real-time PCR. The entire reactions were performed in triplicate (as per MIQE Guidelines) and the total volume of the reaction mixture was set up to 20µl. The reaction mixture is set up with **TYMP/ECGF1, TFAM, TK2** gene and GAPDH (housekeeping gene) primers calculated as per concentration. 1µl of cDNA, 10µl Hi-SYBr Master Mix (HIMEDIA MBT074) and rest volume adjusted with nuclease-free water to achieve the total reaction mixture volume. Then the reaction plate placed into the ABI 7500 system and run the reaction program. The delta-delta-Ct (ΔΔCt) method was used for the analysis of the result. The primers used for the reaction are as followed:

**Table 3:**
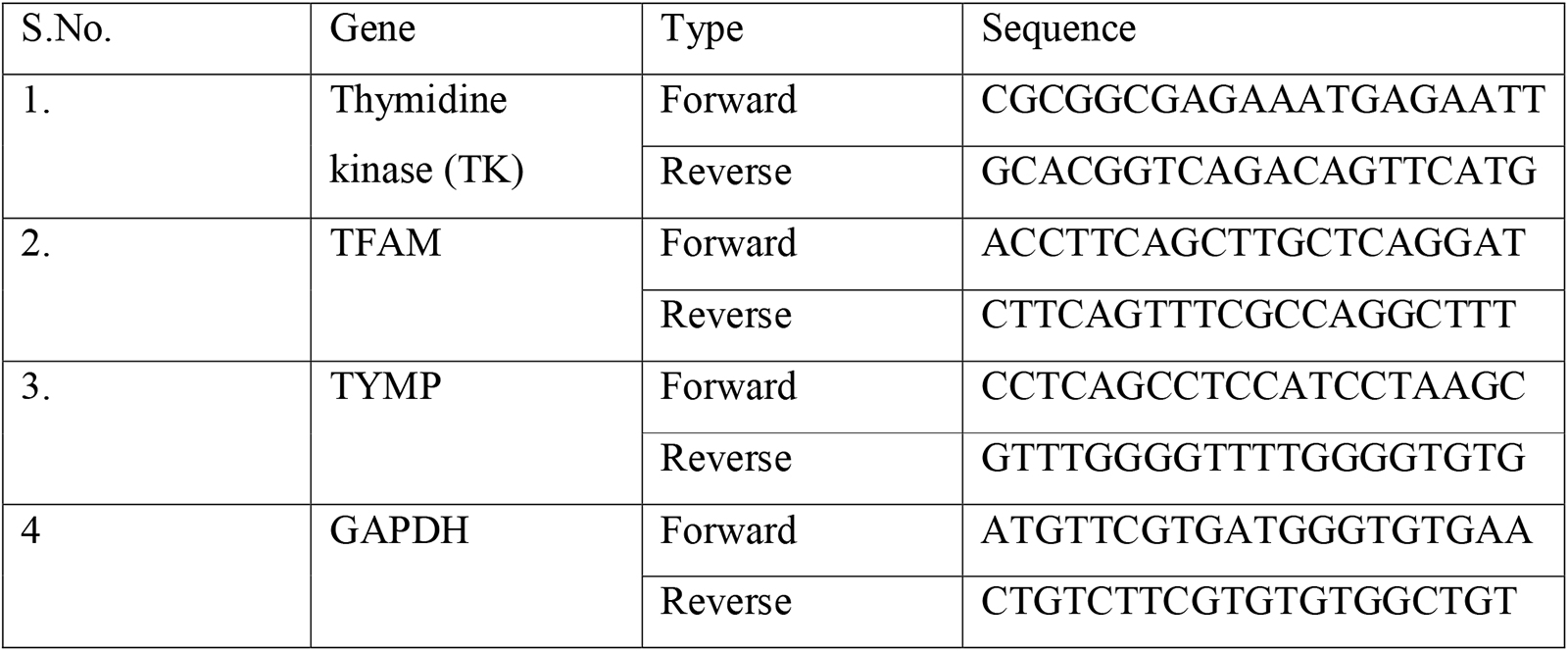
Primers used for the differential mRNA expression profiling of immune response genes of duck. The NUMT genes selected were:

#### Estimation of haematological and biochemical parameters

##### Haematological Profiles

The haematological parameters like haemoglobin, erythrocyte sedimentation rate (ESR) and packed cell volume (PCV) were estimated in whole blood soon after the collection of blood. Haemoglobin was estimated by acid haematin method (Benjamin, 1985), E.S.R. and PCV by Wintrobe’s tube (Hawk, 1965). The total erythrocyte count (TEC), total leukocyte count (TLC) and Differential leukocyte count (DLC) were studied by standard methods described by Jain (1986).

##### Biochemical Analysis

The serum biochemical parameters, estimated in the experiments were total protein, albumin, globulin, albumin: globulin, aspartate aminotransferase (AST), alanine aminotransferase (ALT), alkaline phosphatase (ALP), Total bilirubin, Indirect bilirubin, direct bilirubin, glucose, uric acid, urea and BUN by using a semi-auto biochemistry analyzer (Span diagnostic Ltd.) with standard kits (Trans Asia Bio-Medicals Ltd., Solan, HP, India). The methodology used for the estimation of total protein, albumin, total & direct bilirubin, ALT, ALP, glucose, creatinine urea and uric acid were the biuret method, bromocresol green (BCG) method, 2-4-DNPH method, modified kind and king’s method, GOD/POD method, modified Jaffe’s Kinetic method. GLDH-urease method and trinder peroxidise method respectively.

##### Histological section

The liver samples were fixed in formalin (10%) and embedded in paraffin and processed for histological examination and stereology. The liver tissues were submerged in Lillie fixative for 1 week at room temperature and then were processed and embedded vertically in paraffin wax. Then, each liver sample was exhaustively sectioned into 4 μm-thick sections by a fully automated rotary microtome (Leica RM2255, Germany). Each of these sections was stained with hematoxylin and eosin and mounted. From each liver sample, 10-15 sections were chosen by the systematic uniform random sampling (SURS) method.

##### Statistical Analysis

Microsoft Excel was used for the descriptive statistical analysis. SYSTAT 13.1 software (SYSTAT Software Inc.) was used for statistical analysis and analysis of variance (ANOVA) was used to test between parameters.

## Result

### Characterization of TYMP/ECGF1, TFAM, TK2 gene in Anas platyrynhos

In the current study, four genes are selected, namely Thymidine phosphorylase/ endothelial cell growth factor1, TFAM Transcription factor A, TK2 Thymidine kinase 2 and MPV17. These genes are basically nuclear genes, but coding for mitochondrial protein. This is the first report of such nuclear mitochondrial (NUMT) genes in any livestock or poultry disease.

### Characterization of Thymidine phosphorylase/ endothelial cell growth factor1 gene of duck

The amino acid sequence for Thymidine phosphorylase in *Anas platyrhynchos* is comprised of 464 amino acids. It is derived from the gene sequence. The 3D structure of Thymidine phosphorylase is depicted in Fig 1 in orange color. The thymid phosphorylase (Thymidine and pyrimidine-nucleoside phosphorylases) domain has been depicted at amino acid position **116 – 131.**

**Fig 1:**
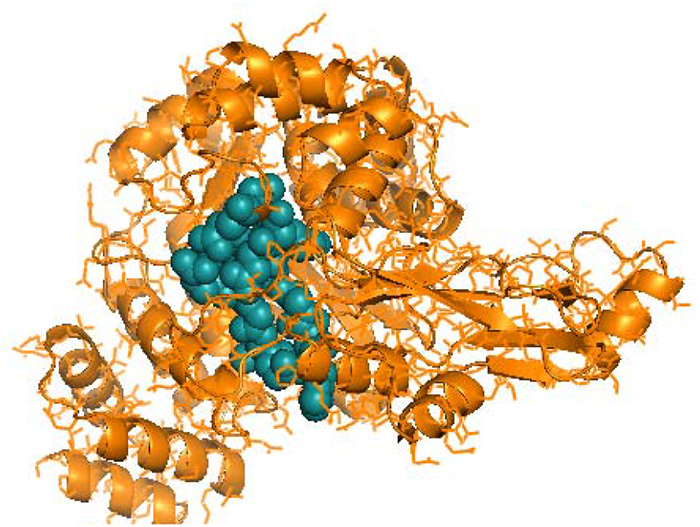
3D structure of Thymidine phosphorylase with THYMID_PHOSPHORYLASE *Thymidine and pyrimidine-nucleoside phosphorylases signature*: at amino acid position 116 – 131 of of *Anas platyrhynchos* .

Thymidine phosphorylase is also known as TdRPase, Gliostatin and Platelet-derived endothelial cell growth factor (PD-ECGF) (Akiyama *et.al*., 2004, NCBI, 2022 Gene ID: 1890)

In mice, it is comprised of 471 amino acids, in rat 476 amino acids and 482 amino acids in humans. (Uniprot, P19971 · TYPH_HUMAN, Uniprot, Q99N42 · TYPH_MOUSE Uniprot, Q5FVR2 · TYPH_RAT)

It has been reported to be involved in mitochondrial genome maintenance. (Nishino *et al*., 1999)

### Characterization of <u>TFAM</u> Transcription factor A, mitochondrial

The amino acid sequence for <u>TFAM</u> Transcription factor A, mitochondrial in *Anas platyrhynchos* is comprised of 272 amino acids. It is derived from the gene sequence. The 3D structure of <u>TFAM</u> Transcription factor A, mitochondrial is depicted in Fig 2 and Fig 3 in red color. HMG Box A DNA binding domain (green, Fig 2) and HMG Box B DNA binding domain (blue, Fig 3) of TFM of *Anas platyrhynchos* have been depicted at amino acid position 45-113 and 157-221 amino acid positions respectively.

**Fig 2:**
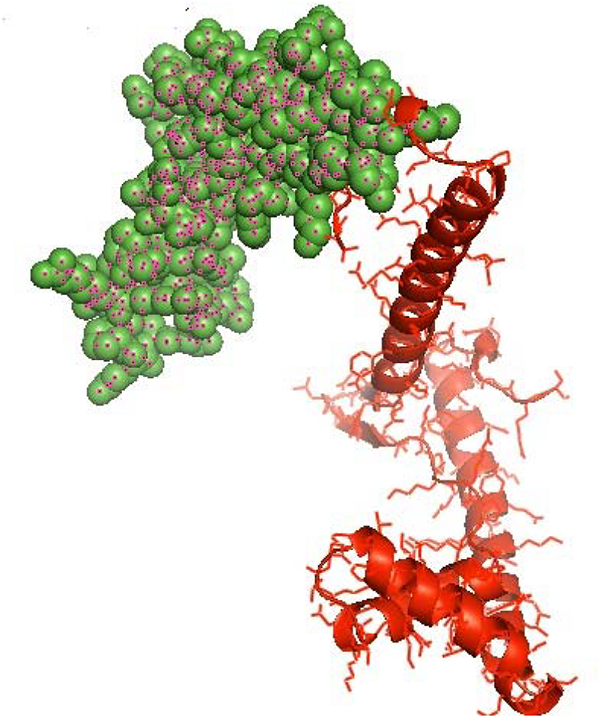
HMG Box A DNA binding domain (green) profile of TFM of *Anas platyrhynchos*.

**Fig 3:**
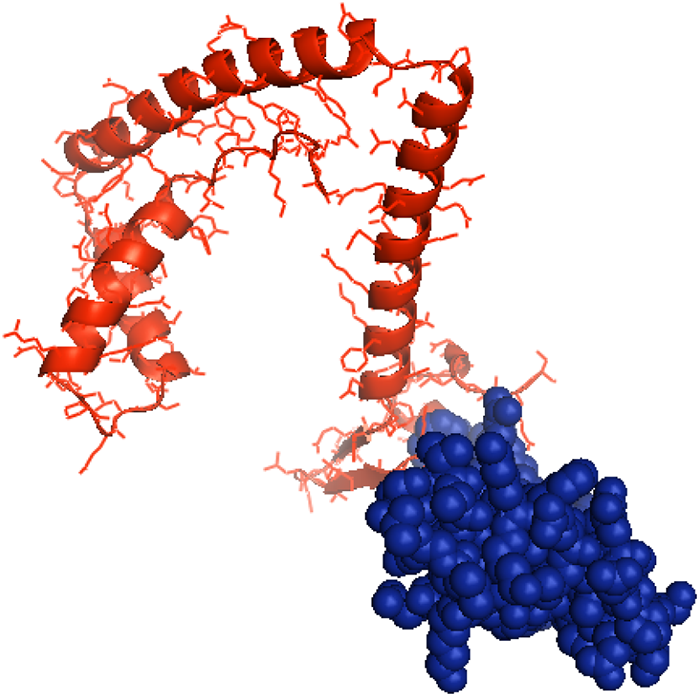
HMG Box B DNA binding domain (blue) profile of TFM of *Anas platyrhynchos*.

TFAM gene is also known as TCF6, TCF6L2, Hmgts mtTFA. (Uniprot, Q00059 · TFAM_HUMAN, Uniprot, P40630 · TFAM_MOUSE, Uniprot, Q91ZW1 • TFAM_RAT)

It is comprised of 246 amino acids in human, pigs and bovine, 244 amino acids in rats, 243 amino acids in mouse. (Uniprot, Q00059 · TFAM_HUMAN, Uniprot, Q0II87 · TFAM_BOVIN, Uniprot, Q5D144 · TFAM_PIG, Uniprot, Q91ZW1 • TFAM_RAT, Uniprot, P40630 • TFAM_MOUSE)

It encodes for proteins Mitochondrial transcription factor 1 (MtTF1), Transcription factor 6 (TCF-6), Transcription factor 6-like 2 in humans, Testis-specific high mobility group protein (TS-HMG) in rats. (Uniprot, Q00059 · TFAM_HUMAN, Uniprot, P40630 • TFAM_MOUSE, Uniprot, Q91ZW1 • TFAM_RAT)

### Characterization of <u>TK2</u> Thymidine kinase 2, mitochondrial

The amino acid sequence for <u>TK2</u> Thymidine kinase 2 in *Anas platyrhynchos* is comprised of 289 amino acids. It is derived from the gene sequence. The 3D structure of Thymidine phosphorylase is depicted in Fig 4 in pink color.

**Fig 4:**
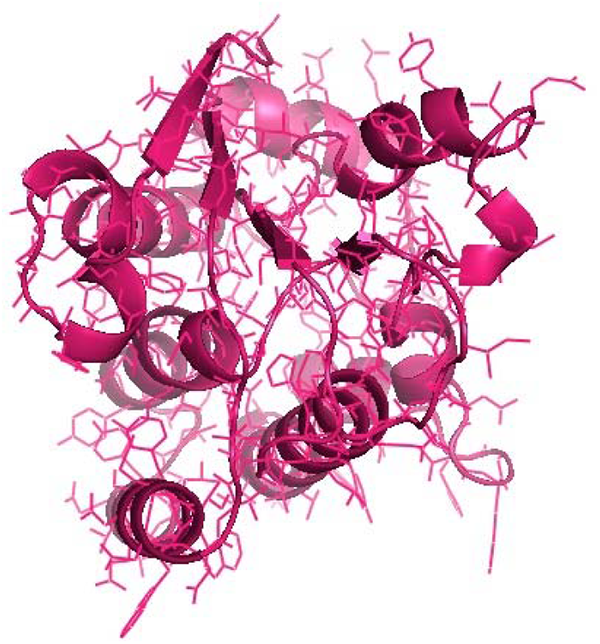
3D structure of Thymidine kinase 2, mitochondrial of *Anas platyrhynchos*.

This gene encodes 265 amino acids in human and *Macaca fascicularis*, 270 amino acids in mouse. (Uniprot, O00142 · KITM_HUMAN, Uniprot Q9N0C5 · KITM_MACFA, Uniprot Q9R088 · KITM_MOUSE)

This gene encodes for prtoteins known as 2’-deoxyuridine kinase TK2, Deoxycytidine kinase TK2, Mt-TK. (Uniprot, O00142 • KITM_HUMAN)

### Differential mRNA expression profiling for NUMT genes with respect to disease resistance against Duck pasteurellosis in *Anas platyrhynchos*

Unvaccinated ducks were naturally challenged with duck Pasteurellosis. The diagnosis was initially conducted through symptomatic diagnosis, followed by final molecular diagnosis of *Pasteurella multocida* with following primer pairs. Individual differences in susceptibility or resistance were observed for Bengal ducks with respect to infection of *Pasteurella multocida*.

NUMT genes as Thymidine phosphorylase/ endothelial cell growth factor1 gene, <u>TFAM</u> Transcription factor A, mitochondrial, <u>TK2</u> Thymidine kinase 2, mitochondrial genes were upregulated in healthy in comparison to infected ducks(Fig 5,6,7):

**Fig. 5:**
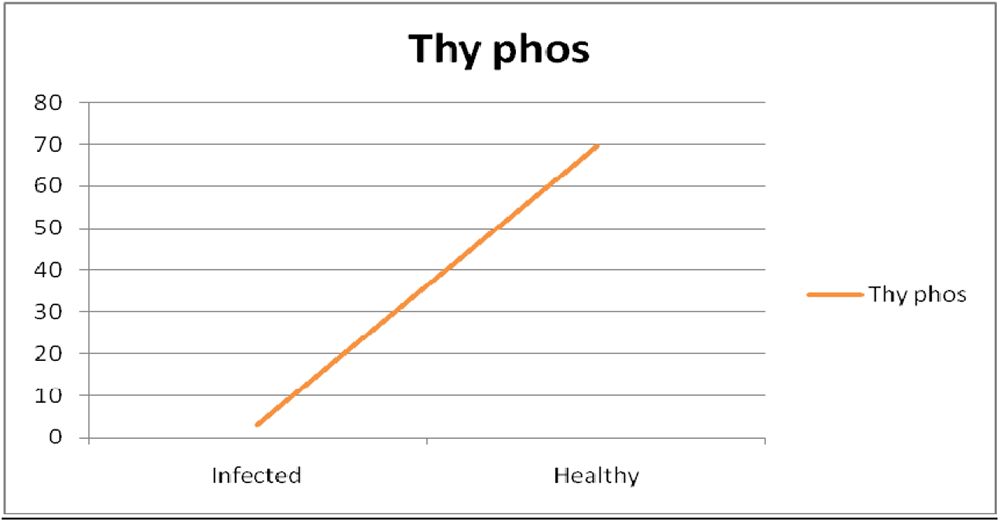
Differential mRNA expression profiling for as Thymidine phosphorylase/endothelial cell growth factor1 gene (NUMT genes) with respect to disease resistance trait in duck.

**Fig. 6:**
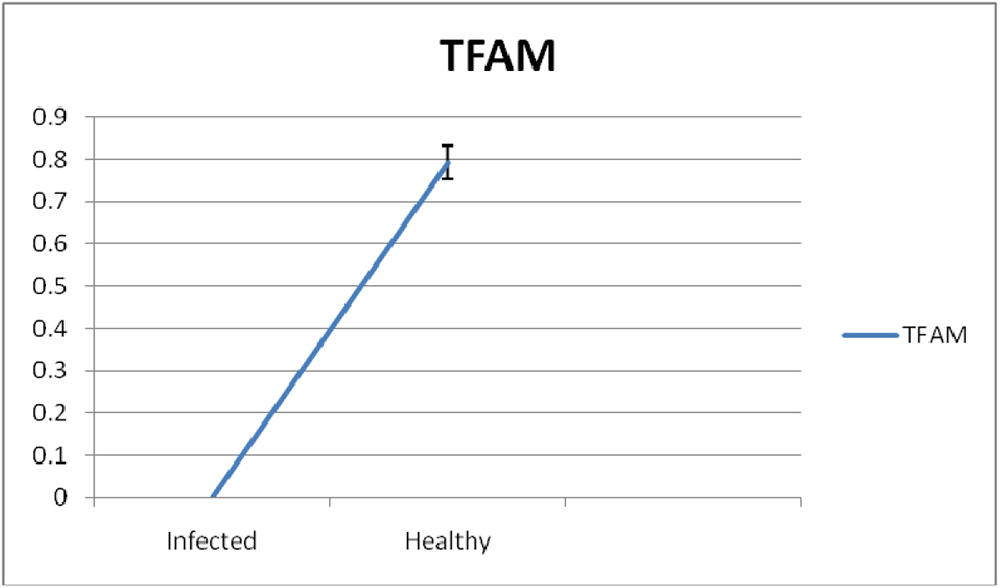
Differential mRNA expression profiling for <u>TFAM</u> Transcription factor A, mitochondrial gene (NUMT genes) with respect to disease resistance trait in duck.

**Fig. 7:**
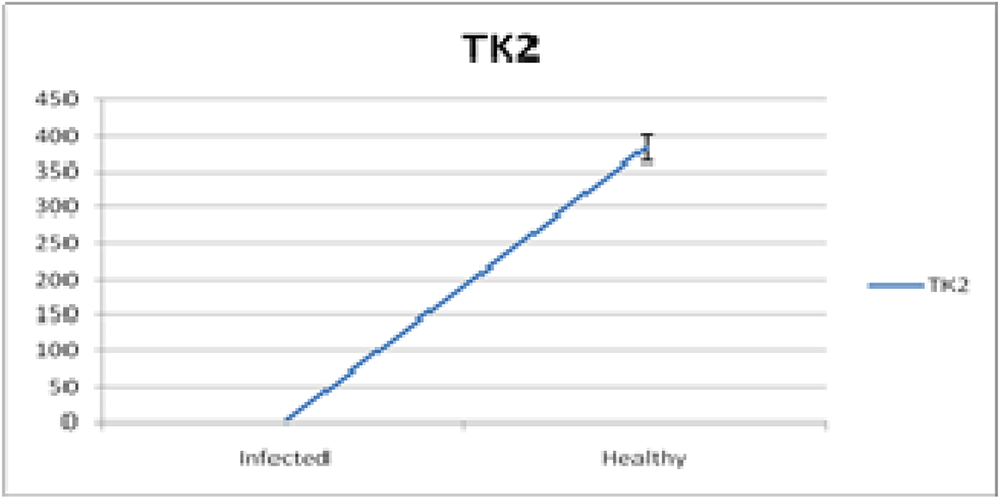
Differential mRNA expression profiling for <u>TK2</u> Thymidine kinase 2, mitochondrial gene (NUMT genes) with respect to disease resistance trait in duck.

## Discussion

Disease resistance is a complex trait controlled by polygenes. We usually designate them as immune response genes, mostly of nuclear origin. In our earlier studies, we could explore certain immune response genes against parasitic infestation in sheep model (reference) and against bacterial (Chakraborty et al., 2023), viral (Pal et al.,2021 and Pal et al., 2020) in chicken and duck model (Pal et al., 2022). Simultaneously, we have also detected certain genes of mitochondrial origin, which are responsible for disease resistance in sheep (Banerjee et al., 2021, Rawat et al., 2021), chicken and duck model (Pal et al., 2020, Pal et al., 2021,Pal et al., 2022). In the following research work, it was attempted to study a novel area of role of certain nuclear genes that act through mitochondria (Chakraborty et al., 2023). It is evident that apart from oxidative phosphorylation, mitochondria plays a varieties of functions including Calcium transporters, Cell proliferation and differentiation, Free radical production, Apoptosis and autophagy, Ageing, Signaling, Immunity, Growth and Reproduction. (Gunter *et al*., 2000, Maeda and Chida, 2013, Cadenas and Davies, 2000, Angajala *et al*., 2018, Pradhan *et al.,* 2018, Pal *et al.,* 2019)

In this current article, we have explored another set of genes which are basically of nuclear origin, but they act through mitochondria, designated as nuclear mitochondrial DNA (NUMT). In healthy animals, there is upregulation of NUMT genes, which means they are expressed better. While in pathological condition or disease condition, there is downregulation of the genes because there is cellular damage, mitochondrial damage, and tissue damage. In healthy animals, upregulation of these genes causes better cell growth, reproduction and metabolism. Upregulation of these genes are responsible for better nerve impulse transmission, mitochondrial and cellular homeostasis, better feed efficiency, and growth. Whereas in pathological condition, there is cellular level stress and hence the genes are not expressed well. The comparison of the disease resistance traits was not possible because it is the first report of NUMT genes on disease traits of ducks.Although we had reported certain preliminary studies on numtogenesis (Banerjee et al., 2020). There are reports that mutation in the mitochondrial gene Cytochrome B causes debility and ill effects on health of sheep. (Pal *et al*., 2019)

Singh *et al*., (2017) reported that Numtogenesis or insertion of the NUMTs into the nuclear genome of somatic cells can alter the pathways contributing to tumorigenesis and disrupt the functions of tumor suppressor genes, leading to development of cancer. The NUMTs sometimes can also directly or indirectly activate oncogenes and thereby further drive development of tumor. Srinivasainagendra *et al*., (2017) in their studies also revealed that Numtogenesis may lead to the development of cancer. Borensztajnm *et al*., (2002) reported that the integration of mtDNA in the IVS 4 acceptor site and human factor VII gene causes deleterious mutation effect, resulting in severe plasma factor VII deficiency and bleeding diathesis. Turner *et al*., (2003) in their studies conducted on a patient with Pallister-Hall syndrome characterized a patient with the de novo nucleic acid transfer from the mitochondrial genome to the nuclear genome, They demonstrated that this transfer or Numtogenesis is a novel mechanism of human inherited disease. Goldin *et al*., (2004) in their studies revealed an inherited transfer of mitochondrial DNA fragment into the nuclear DNA, causing Numtogenesis which leads to genetic disease Mucolipidosis IV. In healthy animals, there is upregulation of NUMT genes, which means they are expressed better while in pathological conditions or disease conditions, there is downregulation of the genes because there is cellular, mitochondrial and tissue damage. In healthy animals, upregulation of these genes causes better cell growth, reproduction and metabolism. Upregulation of these genes is responsible for better nerve impulse transmission, mitochondrial and cellular homeostasis, (Nishino *et al.,* 1999) better feed efficiency and growth. Whereas in a pathological condition, there is cellular level stress and hence the genes are not expressed well.

Nishino *et al.,* (1999) and Gamez *et al.,* (2002) reported that TYMP gene is involved in the disease of Mitochondrial DNA depletion syndrome 1, MNGIE type (MTDPS1) in humans which is a multisystem disease associated with mitochondrial dysfunction. This disease is clinically characterized by its onset between the second and fifth decades of life; there is ptosis, gastrointestinal dysmotility (often pseudoobstruction), progressive external ophthalmoplegia, cachexia, diffuse leukoencephalopathy, peripheral neuropathy and myopathy.

Meyer and Felix (2005) and Muller *et al.,* (1997) reported that mice lacking expression of MPV-17 protein develop adult-onset nephrotic syndrome and chronic renal failure. There was also severe morphological degeneration of the cochlear structures, organ of Corti, of the inner ear along with alterations of the basement membrane of the capillaries. In *Homo sapiens*, depletion of the MPV-17 gene causes Mitochondrial DNA depletion syndrome 6 (MTDPS6), which is a disease due to dysfunction of the mitochondria. It is characterized by infantile onset of progressive liver failure, often leads to death in the first year of life, peripheral neuropathy, acral ulceration and osteomyelitis leading to autoamputation, corneal scarring, cerebral leukoencephalopathy and recurrent metabolic acidosis with intercurrent infections, failure to thrive, neurologic involvement such as delay in development, hypotonia, microcephaly and motor and sensory peripheral neuropathy.

Stiles *et al.,* (2016) reported due to mutation in the TFAM gene, a severe disease condition is seen, known as “Mitochondrial DNA depletion syndrome 15”, hepatocerebral type (MTDPS15), which is characterized by severe intrauterine growth restriction, neonatal-onset hypoglycemia and liver dysfunction, there is also depletion of mitochondrial DNA in the liver and skeletal muscle and abnormal mitochondrial morphology is observed in skeletal muscle. Hepatic pathology includes cirrhosis, steatosis and cholestasis. There is rapid progression to liver failure and death with no neurological impairment or other organ involvement.

Mutation in the Thymidine kinase 2 gene causes the disease “Mitochondrial DNA depletion syndrome 2 (MTDPS2)”, which is characterized by the childhood onset of muscle weakness associated with the depletion of mtDNA in skeletal muscle. There is wide clinical variability; some patients have onset in infancy and show a rapidly progressive course with early death due to respiratory failure, whereas others have a later onset of a slowly progressive myopathy in humans. (Saada *et al.,* 2001, Mancuso *et al.,* 2002, Wang *et al.,* 2005, Tulinius *et al.,* 2005 and Knierim *et al.,* 2015).

## Conclusion

The better expression profile for the other two genes was also observed as TFAM and TK2 affecting reproduction. NUMT genes TYMP, TFAM and TK2 were upregulated in healthy in comparison to infected ducks.

## Acknowledgement

The authors are thankful to Department of Biotechnology, Ministry of Science and Technology, Govt. of India (Grant number BT/PR24310/NER/95/649/2017) and Department of Science and Technology, Govt. of India (Grant no. EMR/2016/003554) for providing the financial support. The technical and financial support by Vice-Chancellor, West Bengal University of Animal and Fishery Sciences is duly acknowledged. Thanks to Director, AH & VS, Animal Resource Development Department, Govt. of West Bengal.

